# LimbNET: collaborative platform for simulating spatial patterns of gene networks in limb development

**DOI:** 10.1101/2024.08.07.607075

**Authors:** Antoni Matyjaszkiewicz, James Sharpe

## Abstract

Successful computational modelling of complex biological phenomena will depend on the seamless sharing of models and hypotheses among researchers of all backgrounds - experimental and theoretical. LimbNET, a new online tool for modelling, simulating and visualising spatiotemporal patterning in limb development, aims to facilitate this process within the limb development community. LimbNET enables remote users to define and simulate arbitrary gene regulatory network (GRN) models of 2D spatiotemporal developmental patterning processes. Researchers can test and compare each others’ hypotheses - GRNs and predicted spatiotemporal patterns - within a common framework. A database of previously created models empowers users to simulate, explore, and extend each others’ work. Spatiotemporally-varying gene expression intensities, derived from image-based data, are mapped into a standardised computational description of limb growth, integrated within our modelling framework. This enables direct comparison not only between datasets but between data and simulation outputs, closing the feedback loop between experiments and simulation via parameter optimisation. All functionality is accessible through a web browser, requiring no special software, and opening the field of image-driven modelling to the full scientific community.

## Introduction

A key challenge for the future of computational modelling is how to easily share models, simulations, and hypotheses between a diverse community of theoreticians and experimentalists. This is essential so that different researchers can test and compare each others’ hypotheses and to allow their new results to build on previous discoveries. In this publication we present LimbNET, a new accessible online simulation tool for intuitive image-driven computational modelling, simulation and visualisation of gene expression patterns important to limb development, and the sharing of associated results within the community.

At the heart of biology is the concept of mechanism. By creating and comparing mechanistic hypotheses and models(Sharpe, 2017) of how something works, we may infer the underlying behaviour of complex biological systems in various model systems, e.g. Drosophila development(Jaeger *et al*, 2004b), neural tube(Balaskas *et al*, 2012) and limb development(Uzkudun *et al*, 2015; Onimaru *et al*, 2016; Raspopovic *et al*, 2014). While data and analysis tools are increasingly shared online, competing hypotheses are still largely the preserve of each individual lab. With LimbNET we propose a new approach: the sharing of hypotheses formalised as mathematical models and concrete computational simulations, integrated with associated experimental data through a user-friendly common framework. In this respect, LimbNET’s core goals are:

- Sharing and accessibility,
- Standardisation of data and models/hypotheses via a common platform: an atlas of 2D spatiotemporal gene expression patterns and a repository of models/simulations,
- Reducing the “energy barrier” of the modelling process, especially for non-experts.

More precisely, our desired system must facilitate investigation of novel signalling networks, integrate models with related data which is also hosted on the same platform (within a common frame of reference), permit online simulation, and collation of models and corresponding inputs/outputs into a unified database that permits model comparison. It should facilitate the exploration of hypotheses - and thus models - both published and unpublished. It should be trivial for a scientist to explore an existing model or, indeed, to challenge it. The platform should support not only tweaking and exploration of models but also more rigorous parameter fitting and optimisation of models onto existing experimental data(Mousavi & Lobo, 2024; Uzkudun *et al*, 2015).

With LimbNET we hope to implement the above goals by addressing the following challenges:

Firstly, within the community of researchers who study limb development, scientists and labs are globally geographically dispersed, and there is a large number of genes, pathways and mechanisms to investigate, necessitating a coordinated approach. Communication is generally indirect and asynchronous (research papers and scientific conferences). In order to integrate and coordinate data and ideas across a geographically diverse research community, LimbNET’s primary interface consists of an accessible web client, central database, and a common server platform. There is no need for scientists to install software, build further modelling tools or compile libraries in order to collaborate on each others’ models(Byrne *et al*, 2010). In the last decade the trend towards online collaboration and web applications has only accelerated; this ongoing proliferation of online platforms has led to a gradual shift in philosophy of data management, analysis, and modelling from predominantly-local, to cloud-based. Precipitated by online (as opposed to physical media(Emmert *et al*, 1994)) databases, the modern web-based science ecosystem now encompasses not only visual analysis tools(Lyons *et al*, 2022) but finally online modelling and/or simulation(Fortmann-Roe, 2014; Wortel & Textor, 2021), with established platforms such as Netlogo(Tisue & Wilensky, 2004) also providing online versions.

Secondly, not only are hypotheses and models geographically spread, but also data. Open online distribution of data has long been recognised as crucial to accelerating bioscience(Emmert *et al*, 1994; Baxevanis & Bateman, 2015). While publications disseminate results, this does not always guarantee or ensure access to data. Although freedom of - and ease of access to - experimental data is rapidly increasing, even in the case that data are shared, non-standardised interchange formats may complicate re-use. LimbNET’s centralised database of gene expression patterns from the limb development community, along with mapping and interactive exploration aims to rectify this by providing researchers with an atlas of genes in 2D and time for (eventually all) gene expression patterns relevant to limb development. It is stored in such a way that all hypotheses/models can access all data in the same standardised manner.

The field of limb development is rich with data; many aspects of limb (bud) development are well-characterised, and it is a paradigm model system of the more generalised problem of developmental patterning and organogenesis(Petit *et al*, 2017; Capdevila & Belmonte, 2001). The limb bud exhibits a rich variety of dynamical patterning behaviours during development(Benazet & Zeller, 2009; Raspopovic *et al*, 2014). Of these, the vast majority of characterised behaviours have been described primarily experimentally. The developmental trajectory of the limb bud consists of a well-defined time course, and so data can be accurately staged and mapped into a consistent common reference framework(Musy *et al*, 2018). Within LimbNET we provide tools for staging and digitisation that are tied into the database and model framework. This allows provenance to be tracked, back to the original imaging data which can finally be used to fit model parameters, closing the modelling loop.

The final challenge - and the hardest - is that of mixed expertise throughout the field. Existing modelling and simulation frameworks tend to be generalised(Mirams *et al*, 2013; Starruß *et al*, 2014), requiring significant effort and domain-specific knowledge in order to create a general model of limb development. Some classes of models can also be shared through centralised repositories such as BioModels(Malik-Sheriff *et al*, 2020), or as part of the community for a specific tool as in the case of Morpheus(Starruß *et al*, 2014). However, simulation and modelling is limited to a small group of specialised “theoreticians”, while our goal is to bring experimental researchers into this endeavour. While researchers may not be familiar with the in-depth details required to implement a mathematical model, they should nonetheless be able to leverage an existing framework to simulate their hypotheses, thus concretising abstract ideas. Our core goal is to be able to rapidly gain intuition of a given system’s behaviour, to tighten the iterative loop(Harline *et al*, 2021) of model design, building and testing. Indeed, while it is common to study genes one by one, disentangling the complex causal relationships of a system’s network dynamics requires a systems biology approach. By providing a straightforward interface for definition and simulation of models, we hope to lower the “energy barrier” for model development and streamline the modelling process. Our repository of models/simulations - including examples, tutorials and published hypotheses - will facilitate creation of models, either *de novo* or by extending/modifying an existing model/hypothesis leading to a truly integrated and collaborative modelling process where scientists build on others’ work, while testing and comparing existing hypotheses. Finally we hope that such a system may provide an improved ontology, a platform upon which to build standardisation of limb development terminology (e.g., a controlled and well specified coordinate system).

Only a minimal proportion of publications contain a significant theoretical/modelling perspective, thus there exists a large quantity of unexploited gene expression data, primarily in 2D, that has been published over the last decades yet has not been used in mathematical model development. Clearly, a profound range of tested hypotheses are associated with these data, predicting interactions that lead to patterning behaviours, however we believe that by formalising these hypotheses as mathematical models we can gain further insight into the predicted mechanisms. The collection of researchers studying limb development is a tightly-knit community, and therefore creating a tool that can be used by the whole community is a tractable problem, permitting development of a useful tool that provides sufficiently specific functionality while nonetheless remaining general enough to be used in other applications.

It is crucial to note that while many generalised modelling platforms already exist, general-purpose simulation tools may sometimes cover too broad a scope, leading to a proliferation of different tools as users try to customise existing solutions or develop their own to suit their own modelling pipelines. In this study we instead focus on high quality modelling of a specific system: patterning GRN models on a well-defined growing domain of limb development, in two dimensions.

### Implementation

LimbNET’s design follows a client-server architecture (Figure 1). There is a direct logical separation of the user-facing interface, referred to as the LimbNET client (Figure 1A), and the underlying implementation of models (Figure 1B) and simulations/data storage, referred to as the server or backend (Figure 1C). For the user - typically an experimental scientist - this has numerous advantages: for instance there is no need to deal with problems of data representation, storage, or computation, all of which are dealt with remotely. The web-based client further simplifies access. Non-specialist users can access all functionality - data visualisation, modelling, and simulation (Figure 1A) - through the web browser without the need for installation (let alone compilation from source).

**Figure 1:**
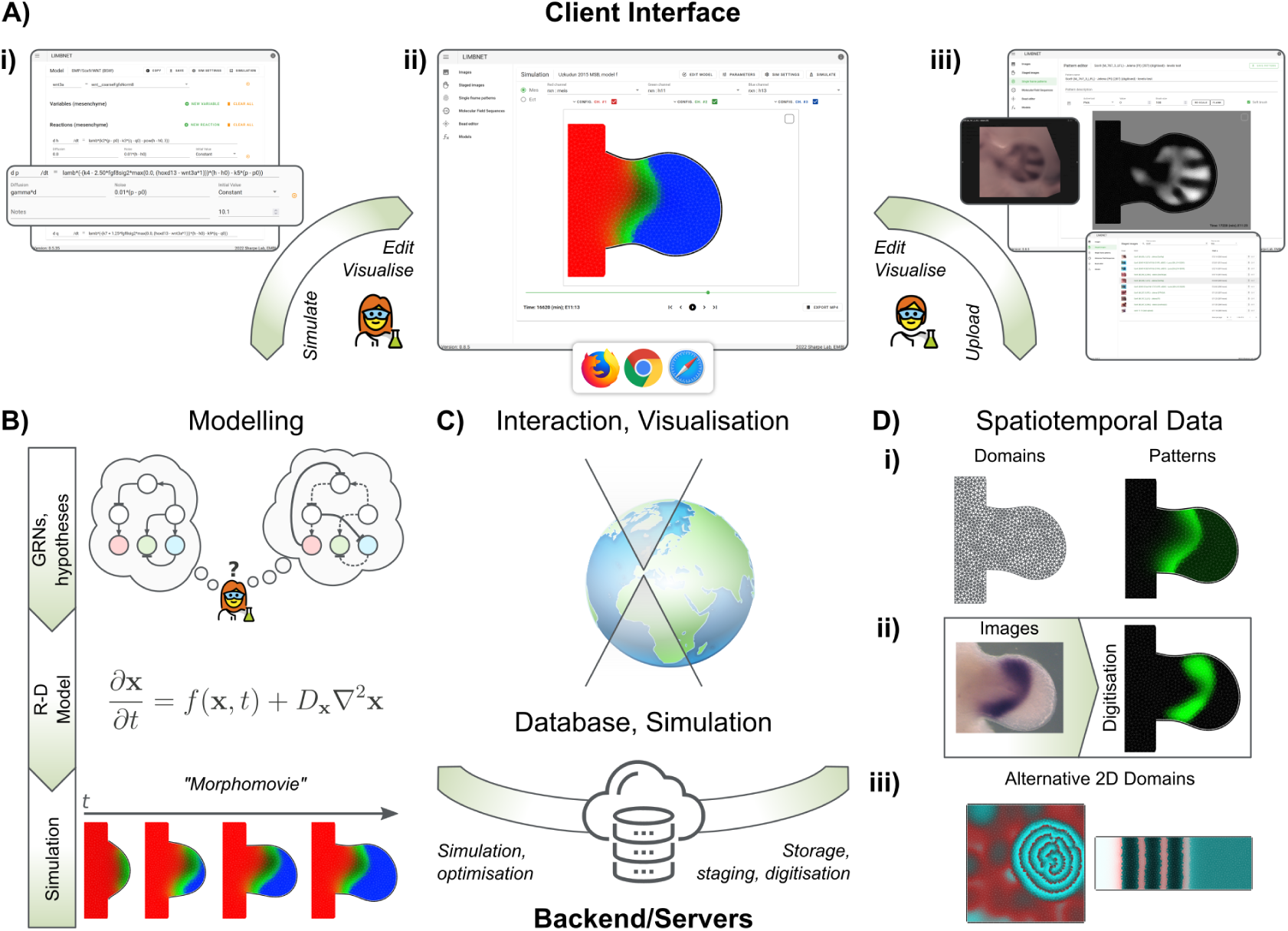
High-level schematic overview of the LimbNET platform. A) LimbNET’s user-facing client is a single-page web application, a unified GUI exposing specific views corresponding to defined modelling or data visualisation tasks: (i) model definition; (ii) visualisation of model simulation; (iii) upload/visualisation of image-based spatiotemporal data, and corresponding digitisation. B) LimbNET facilitates a streamlined workflow for modelling, simulating and comparing patterning hypotheses. Users input GRNs represented by spatial patterning GRN equations ((A,i), above). LimbNET simulates the model, outputting “morphomovies” that illustrate simulated spatiotemporal patterns over a dynamic 2D domain. C) The client GUI communicates with the backend (databases, model simulation engine). Users need not deal with the computational requirements of heavy numerical simulation, nor associated storage of results, which are streamed to the web client as required. D) LimbNET specialises in 2D spatiotemporal data, capturing time-dependent scalar quantities (gene expression, ligand concentration, etc.) over a dynamic 2D domain. (i) Spatiotemporal patterning data are stored internally along with the relevant domain shape(s) - e.g., mouse embryonic hindlimb bud - and their corresponding discretisation (a triangular mesh). (ii) Uploaded image-based data (e.g., *in-situ* hybridization gene expression), stored in our database, can be digitised. Digitisation maps image intensity onto a discretized 2D domain. These patterns can then be used in models and simulations in LimbNET. (iii) Versatility - LimbNET is not limited to limbs, providing alternative pre-defined growing or static meshed domains. (*Left*) a snapshot of the classic “Brusselator”(Prigogine & Lefever, 1968) on a square domain; (*right*) the Progressive Oscillatory Reaction-Diffusion (PORD(Cotterell *et al*, 2015)) model in a rectangular domain.

All relevant functionality is presented as a user-friendly Graphical User Interface (GUI), with no need for prior programming knowledge. It is worth noting that this form of interaction may be disadvantageous for power users; however should the need arise, for example for automation purposes, an Application Programming Interference (API) is also available which may be leveraged programmatically via external scripts and tools. LimbNET has been written using standard technologies that are in widespread use throughout the software industry, and have been proven to behave robustly. Documentation is provided online specifying common usage and tasks, tutorials and examples, and implementation details.

### Client

LimbNET’s client, the core of the user-facing experience, is written in a mixture of javascript and typescript and implemented as a single-page web application (SPA), primarily using the Vue.js framework^1^. The SPA format presents the user with a single interface, similar to a desktop application, with multiple “views” corresponding to different tasks the user may want to perform (Figure 1A): for example, definition of a model specification and corresponding equations (Figure 1A, i); visualisation of spatiotemporal patterns either arising from digitised data or from model simulation outputs (Figure 1A, ii); visualisation, editing and digitisation of experimentally acquired imaging data (Figure 1A, iii).

The application offers task- or data-oriented views, in a hierarchy of searchable and filterable listings for each data type. Specific views for each data type allow users to view/visualise a given dataset, and perform specific data-oriented tasks such as editing related metadata or using data for further experiments. For a given type of data, the owner may decide whether to make it public (available to the whole LimbNET community) or keep it private (viewable only by the owner).

#### Modelling

The core objective of LimbNET is to facilitate the straightforward modelling, simulation and comparison of hypotheses for a given patterning mechanism (Figure 1B). Users input their desired GRN (Figure 2A) to LimbNET as a model specification (Figure 2C). Or indeed users can simply browse and run models which are already uploaded into the system. For example, the model shown in Figure 2 can be found in the web client’s model menu with the name “Uzkudun et al. 2015, model C”. Models represent high-level abstractions of patterning, defined by dynamical systems of coupled reaction-diffusion partial differential equations (PDEs) (Figure 2B). Simulation of a given model produces a single compound output - a “morphomovie” - a collection of spatiotemporal patterns, over a moving and growing 2D domain (Figure 2D). Simulations are thus concrete realisations of a model - a given spatiotemporal patterning process - arising from numerical simulations of a model’s equations under specific parameter sets and virtual experimental conditions.

**Figure 2:**
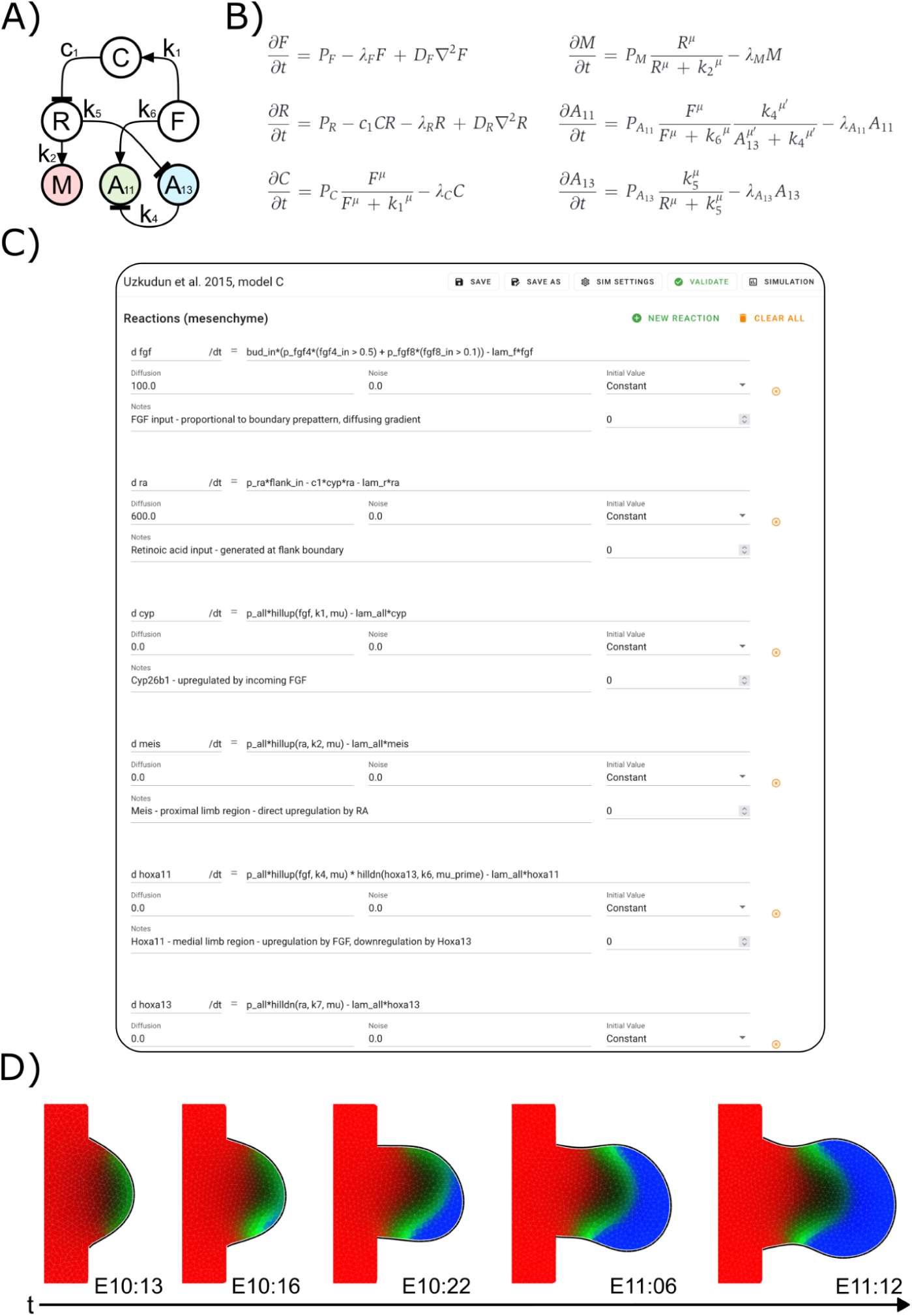
Limb patterning network model specification and simulation. A) One way of describing patterning processes is as a network of interactions - mutual regulation of “morphogens”, genes, metabolites and other relevant spatiotemporal quantities. The network shown describes a possible mechanism for proximo-distal patterning in the limb bud (“Model C”, taken from(Uzkudun *et al*, 2015)); Variables are defined as follows: F, FGF; R, retinoic acid; C, Cyp26b1; M, Meis; A11, Hoxa11; A13, Hoxa13. B) The system of coupled Reaction-Diffusion PDEs corresponding to the network in (A). C) LimbNET’s client-side model editing interface, showing the equations corresponding to the system in (B), as entered by the user, as well as initial conditions, diffusion, and noise where relevant. Additionally the editor allows users to create named variables, global parameters, predefined patterns. Many models such as this one are provided in LimbNET’s model menu, so users do not have to type these from scratch. D) LimbNET’s predicted patterns for Meis (red), Hoxa11 (green) and Hoxa13 (blue) genes, resulting from numerical simulation of the model defined in previous panels.

Users can browse, filter and view all publicly available models and corresponding simulation results, including those from previously published works, tutorials, examples/case studies, and user-shared public models. Users may create *de novo* model specifications, and also copy or fork existing models, in order to further edit them and build upon previous work. Through the model editor (Figure 2C) users may create and freely edit model specifications, and define global parameters, variables, and PDEs. All relevant editable components of a model specification are exposed as freely editable building blocks of the corresponding PDE system. Global parameters and variables can be defined, along with expressions for their respective values. Potential errors and model inconsistencies are highlighted in the GUI.

The core limb bud model consists of two compartments: the mesenchyme (a 2D domain on which the majority of patterning occurs) and an optional ectoderm (the 1D linear boundary of the limb bud). Within compartments, users may freely define and tailor variables and reactions. PDEs are specified according to their derivative’s right-hand-side, along with expressions for diffusion terms, noise and initial conditions. Moreover, users can choose from various geometries for the model/simulation domain (Figure 1D, iii).

In addition to standard variables/reactions, users can specify dynamic “predefined” spatiotemporal pattern variables that evolve over time independently of other model entities. These represent temporal sequences of digitised images, mimicking experimental data for direct comparison of model outputs to experimental observations. Secondarily, they can be chosen to act as feed-forward inputs to the rest of the system, receiving no feedback from other variables. For example, in the model shown in Figure 2, the region where Fgfs are expressed is considered a predefined input to the system, labelled as fgf4_in and fgf8_in. This permits auxiliary functionality such as pre-computing morphogen patterns known to act only as one-way inputs, or defining other experimental interventions such as mobile ligand-soaked beads(Mercader *et al*, 2000).

Upon defining the model, users may initiate simulations under a given domain shape, a set of parameters and a set of inputs. The platform performs the numerical simulation for the given model specification remotely on the server - this way the efficiency of the calculation does not depend on the user’s own computer. The results of the simulation are a set of spatiotemporal patterns which are automatically transferred to the client browser for browsing, comparison and visualisation in the simulation view (Figure 1A, ii). Here the user may view the 2D domain and analyse all the spatiotemporal patterns (3 may be seen at any one time represented by red, green and blue channels in the visualisation). Crucially, the user has the flexibility to freely slide back and forth through time, while adjusting the intensity of each channel independently to visualise the temporal dynamics of their system within the growing and deforming mesh which represents the tissue movements of the limb bud.

#### Image-based data

The simulations create predictions of dynamic gene expression patterns over time (as well as other molecules, such as the retinoic acid in Model C above). But in addition to predicting patterns, it is important to be able to upload known gene expression patterns from data, which generally means image data of *in situ* hybridisations.

In LimbNET users are able to upload and view single-channel images of gene expression patterns in limb buds. Images are generic and may represent qualitative or quantitative acquisition (Figure 1D) including whole-mount *in-situ* hybridization, fluorescence microscopy, *in-situ* hybridization chain reaction (HCR), or other techniques. Our image database can be searched and filtered to find expression patterns related to a specific gene or experimental methodology, for example. Images uploaded and digitised by a user may be kept private for the user, or made public to the community.

#### Staging and digitisation

Image processing functionalities are also provided to users within the web-based platform. LimbNET’s image viewer is integrated with our quantitative Embryonic Mouse Ontogenetic Staging System(Musy *et al*, 2018). Given an image of an embryonic limb the user draws an approximate outline and the system will return the developmental stage of the embryo, based on the morphometry of that limb bud (generally to an error of +/− 1 hour), as well as the canonical outline shape for a limb of that developmental stage (Figure 1D, i).

Staging an image maps it to a canonical limb bud shape and the representation for that given developmental stage. Intensity of a spatial pattern (e.g., gene expression) within the image may then be “digitised”, i.e., mapped onto the canonical domain shape (Figure 1D, ii). A series of digitised patterns, corresponding to ordered time points, may be combined into sequences which may then be used in the modelling and simulation pipeline.

### Server/Backend

LimbNET’s backend - server components - provides the web API, database and all associated computational capabilities needed by the web client. The backend consists of a HTTP server, API server, database, task queues and a computational server. This implementation separates web service functionality - direct communication with clients - from longer-running asynchronous tasks (analysis and computation).

The LimbNET API server is written in python using the Flask^2^ framework, and the database is implemented using PostgreSQL^3^. In order to separate web services from longer running asynchronous jobs we implemented a task queue using Redis^4^ and RQ^5^. Aside from benefitting from a separation of concerns, this makes the simulation system scalable should the need arise: computational tasks can be distributed as we see fit across one or more remote workers, for example during parameter optimisation.

Computational tasks consist of two main categories: analysis of image-based data, and simulation. Image analysis includes staging - as discussed above - which leverages the system published in(Musy *et al*, 2018). Associated digitisation and offline graphical tasks are handled using the Vedo library(Musy *et al*, 2022). Simulation tasks consist of a hierarchy of job dependencies. Firstly, the model specification is validated and parsed. The parsed model specification is then used to generate an internal model representation for our custom simulation engine written in c++. This is the final artefact required to run a simulation job: this task is dispatched to a remote worker, which runs the simulation, reports progress to the client and finally returns a computed result that is stored with the associated model and can be further analysed by the user.

### Simulation

#### Reaction-Diffusion expressions

Simulation in LimbNET consists of the numerical integration of a set of spatial patterning GRN PDEs by our simulation engine, according to a user-supplied model specification (Figure 2). A network of interactions (Figure 2A) represents the interplay of regulatory components that contribute to a patterning process. Our model specification structure represents a system of coupled (stochastic) PDEs (Figure 2B), modelling interactions in time and space over a growing 2D domain. The equations may be expressed in the general form

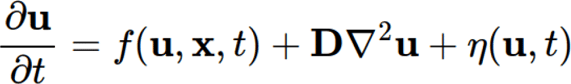

where **u** is a vector of all reactants in the system, *t* is time, **x** is our spatial coordinate, **D** is a diagonal matrix of all per-species diffusivities and η is a vector of all per-species noise terms.

LimbNET’s principal model implements a dynamic 2D domain representing a growing limb bud, which is an extension of the model published in (Marcon *et al*, 2011). The model comprises three compartments: the 2D limb domain (mesenchyme), its 1D boundary (ectoderm), and a non-spatial global compartment. Users may define named parameters/numerical constants as well as time-dependent variables and reactants, either globally or in a spatial compartment.

Reactants in both the ectoderm and the mesenchyme are user-defined equations expressing the reaction component in the right-hand-side of a spatiotemporal derivative - *f*(**u**, **x**, *t*) in the system above (Figure 2C). They may be explicit functions of time, of global entities, and of entities in the same compartment. They are (optionally) subject to diffusion and noise, and are explicitly integrated in time. Diffusion and noise coefficients may additionally be defined as spatiotemporally varying expressions of the same form as compartmental variables.

Compartmental variables are similar to reactants. However - unlike reactants - they are not the right hand side of a derivative, and thus are not integrated in time, neither are they able to diffuse or to be affected by noise. Both variables and reactants have explicit spatial dimensionality corresponding to their compartment, and connections between the mesenchyme and ectoderm compartments are defined in terms of a diffusive coupling between a user-selected pair of reactants.

Furthermore, predefined patterns can be created within each compartment. These may change in space and time, but are fixed with respect to the rest of the model, acting only as feed-forward inputs to the system.

The user may specify as many entities as desired (within practical limits; too many variables may have a prohibitive impact on computation time). The client GUI, exposes the right hand side(s) of the above PDEs as editable fields (Figure 2B), such that *f*(**u**,**x**,*t*), **D**, and η can all be separately written as free mathematical expressions, functions of other user-defined entities in the model. A selection of standard mathematical functions are available for use in all expressions, as well as predefined functions useful for constructing models of GRNs (e.g., Hill-type up-/down-regulation of a reactant). It is up to the user how they define their model: there is no specific restriction to GRNs, nor do variables and reactants in the system need to represent genes; a variable or reactant simply represents any spatiotemporally varying scalar quantity.

#### Simulation domain and spatial discretisation

LimbNET’s simulation engine is designed to model dynamical patterning processes in 2D during early embryonic limb development. The boundary and 2D geometry of the simulation domain is dynamic over time, representing the known tissue-movements during limb development. The standard wildtype growth has been pre-specified, based on analysis of clonal experiments (Marcon *et al*, 2011) - it does not dynamically respond to control from the GRN. While this might be seen as a limitation to modelling certain limb development phenotypes, the current focus of LimbNET is to address the complexities of GRN dynamics *per se*. A large proportion of reported perturbations (genetic and also non-genetic, such as bead experiments) reveals shifts in gene expression patterns without significant growth defects. Indeed, the most powerful perturbation results for the task of deciphering GRNs are this large number which primarily alter molecular patterning. In the future LimbNET will be extended to address growth defects as well, a topic that is addressed further in the discussion.

Using a standard wildtype description of growth also facilitates comparison between models, data, and patterning in general, since it ensures that all spatiotemporal patterns share a common discretized domain. Additionally, since meshing a discretized domain is only precomputed once, the overall speed of computation is improved, especially when it may be necessary to run hundreds of simulations (for example during parameter optimisation). Growth, stretching, and changes in shape are essential to our model, so ensuring the cost of remeshing is only paid once in advance is valuable in order to maintain good performance. Creating the pre-defined dynamical mesh was done as follows. First the boundary was defined, in this case a curve defining the canonical mouse hindlimb developmental trajectory(Musy *et al*, 2018), captured at hourly time points (Figure 3A). This boundary was combined with a small rectangular region corresponding to the embryonic flank, and the entire domain was then discretized as a mesh composed of triangular elements, representative examples of which can be seen in Figure 3B. Additionally, the boundary itself was discretized using linear elements, where any linear element coincides one-to-one with the neighbouring edge of its corresponding triangular element (Figure 3C). Thus the inner domain, triangle-meshed, forms a compartment which effectively represents the limb bud mesenchyme and the outer boundary, formed of linear elements, the corresponding limb bud ectoderm. Material can be exchanged between the mesenchymal and ectodermal compartments, akin to a dynamical boundary condition, which could represent diffusion of a substance between the two compartments.

**Figure 3:**
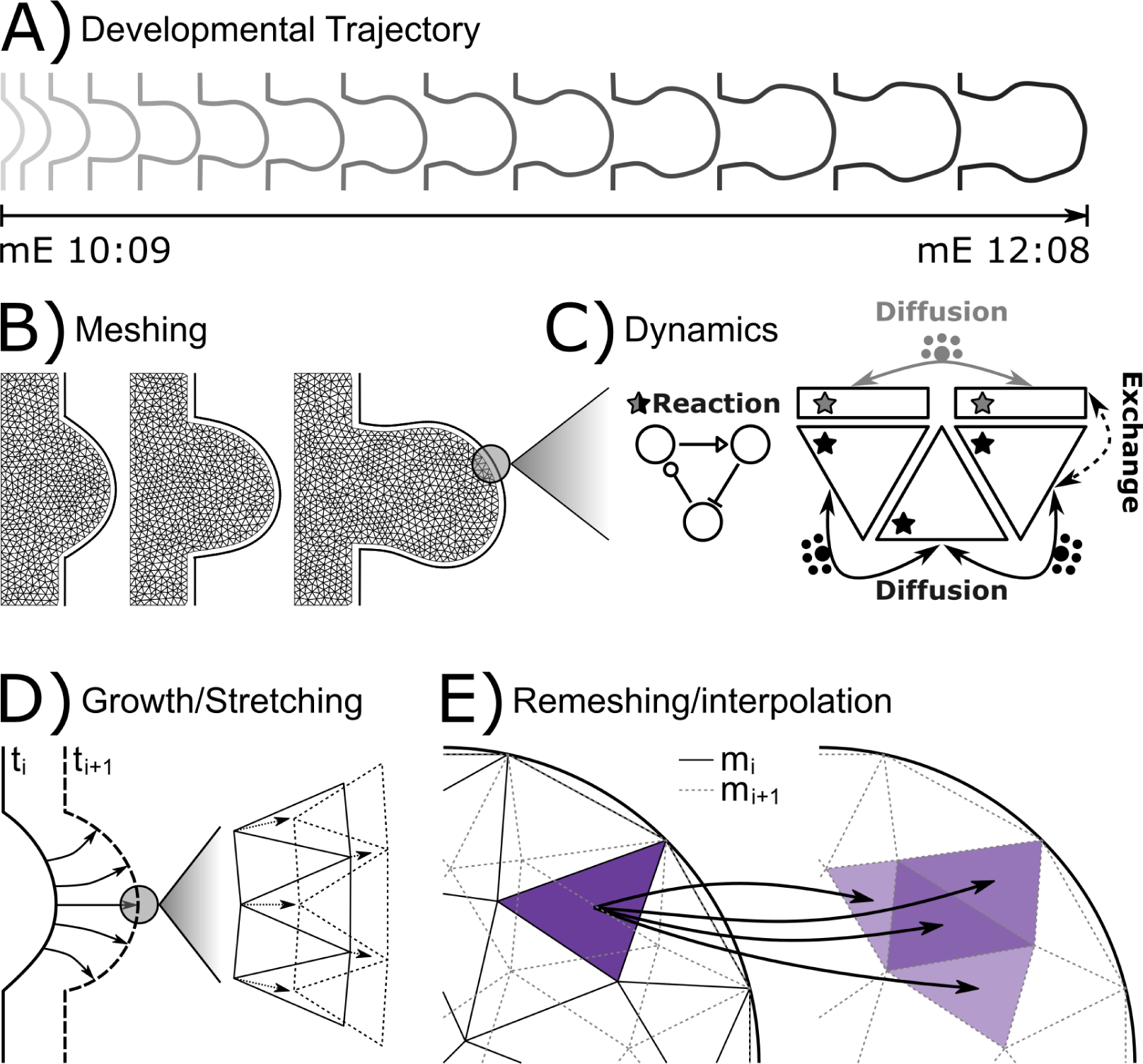
Embryonic limb bud model dynamics and numerical simulation. A) Selected snapshots of canonical mouse hindlimb developmental trajectory(Musy *et al*, 2018) from mouse embryonic stage mE10:09 (10 days and 9 hours), through to mE12:08. Simulated reaction-diffusion processes take place on a 2D domain bounded by this canonical shape trajectory. B) The limb bud boundary (A), with the addition of a rectangular domain representing the flank, is meshed with triangular elements at hourly intervals to form the discretized mesenchyme domain(s) used for subsequent numerical simulations. Three representative meshes are shown; the full trajectory comprises 48 meshes, one per hour of the two-day developmental trajectory. The limb bud boundary (ectoderm; *solid line*) is separately discretized with linear elements. C) Numerical simulation of a model’s GRN reactions. For a single time-step of a given simulation, diffusion is integrated according to a finite volume scheme. In the mesenchyme, diffusion occurs only between neighbouring triangular elements (*black arrows*); in the ectodermal boundary, diffusion occurs only between neighbouring linear elements (*grey arrows*). Material can be exchanged between ectoderm and mesenchyme (*dashed line*). At every time step, reaction proceeds according to the numerical solution of the system of PDEs corresponding to all given GRN species, integrated per individual element (*stars*) in mesenchymal and/or ectodermal compartments as relevant. D) Within each hourly interval a single mesh is defined, however growth and stretching of the simulation domain is treated as a smooth and continuous process. Thus within a given hour of simulated development, the meshed domain (*inset*) deforms such that its original shape (*solid lines*) - derived from the canonical limb bud boundary at the beginning of the hour - fits the boundary at the end of the hour (*dashed lines*)(Marcon *et al*, 2011). E) The mesh is replaced at hourly time intervals, as described in (B). Upon mesh replacement, all per-element scalar quantities (reactants and variables) are interpolated from the mesh at the end of the previous time point (m_i_, *black solid lines*) onto the mesh at the current time point (m_i+1_, *grey dashed lines*). For any given element in the original mesh, a scalar quantity (*purple*) is interpolated onto elements in the new mesh with which it intersects proportionally according to the ratio of overlapping area between the elements.

Whenever the user performs a simulation, numerical integration is performed at every integration time point to solve the PDEs of the system (Figure 3C), for both the reaction and diffusion terms. A combination of an explicit Euler-Maruyama(Kloeden & Platen, 1992) scheme for the reaction terms, and a finite-volume scheme for diffusion is used by the solver. Diffusion is integrated between pairs of neighbouring elements of the same type. Finally, where specified, a diffusion-like exchange of material is computed between neighbouring elements of different types (in this case the boundary elements and those on the interior of the domain). Noise is approximated according to a Wiener process (a one dimensional Brownian motion) scaled per species. The outputs of the resulting simulation can be viewed within the user interface (Figure 2D).

Growth and thus a gradual reshaping of the domain boundary necessitates regular remeshing to avoid degenerate or distorted elements. As in previous studies(Marcon *et al*, 2011; Uzkudun *et al*, 2015; Raspopovic *et al*, 2014) we chose to generate a new mesh at every hourly time-point, corresponding to the time points at which new boundaries are defined. Within each hourly stage, the boundary is assumed to grow smoothly (Figure 3D) and as it grows it induces a stretching and deformation of the corresponding mesh. Finally at the hourly time points where a new mesh has been computed, spatial data from the current time point’s mesh is interpolated onto the next (overlapping) mesh proportional to the amount of overlap between elements in the two meshes (Figure 3E). The stretching within an hourly time interval is defined so that, at the mesh changeover point, the mesh from the previous time interval has deformed such that its boundary overlaps exactly with the new boundary; all material within the domain is conserved.

The simulation framework described here can be generic, and could be applied to other growing and deforming domains (of other organs), but in the LimbNET project we currently focus on the developing limb bud. Extending the system beyond the limb bud would only require new domains, discretisations, and corresponding data structures. Indeed we already define multiple versions of the mouse limb bud with different discretisations (to validate stability of numerical experiments), as well as domains with more basic geometry (used to approximate patterning in cell culture experiments, for example).

### Case Studies

To demonstrate the flexibility of the LimbNET platform in developing different types of models, we implemented two very different previously-published models of patterning processes within the embryonic mouse limb bud.

The first model we chose to replicate was the proximo-distal (PD) patterning model of Uzkudun et al.(Uzkudun *et al*, 2015), a good example of a GRN model to explain dynamic gene expression patterns. From a high-level point of view, the network (as in Figures 2A and 4A) consists of two inputs - retinoic acid (RA), and FGF - representing signals from, respectively, the proximal and distal ends of the growing limb bud. The network is largely composed of feedforward interactions with one feedback, and the regulatory interactions comprise both inhibition and upregulation, modelled using a combination of Hill-type dynamics as well as more general mathematical functions (Figure 2B, 2C). The network was implemented in LimbNET and simulated, obtaining results that agree with those previously published (Figure 4A, i).

**Figure 4:**
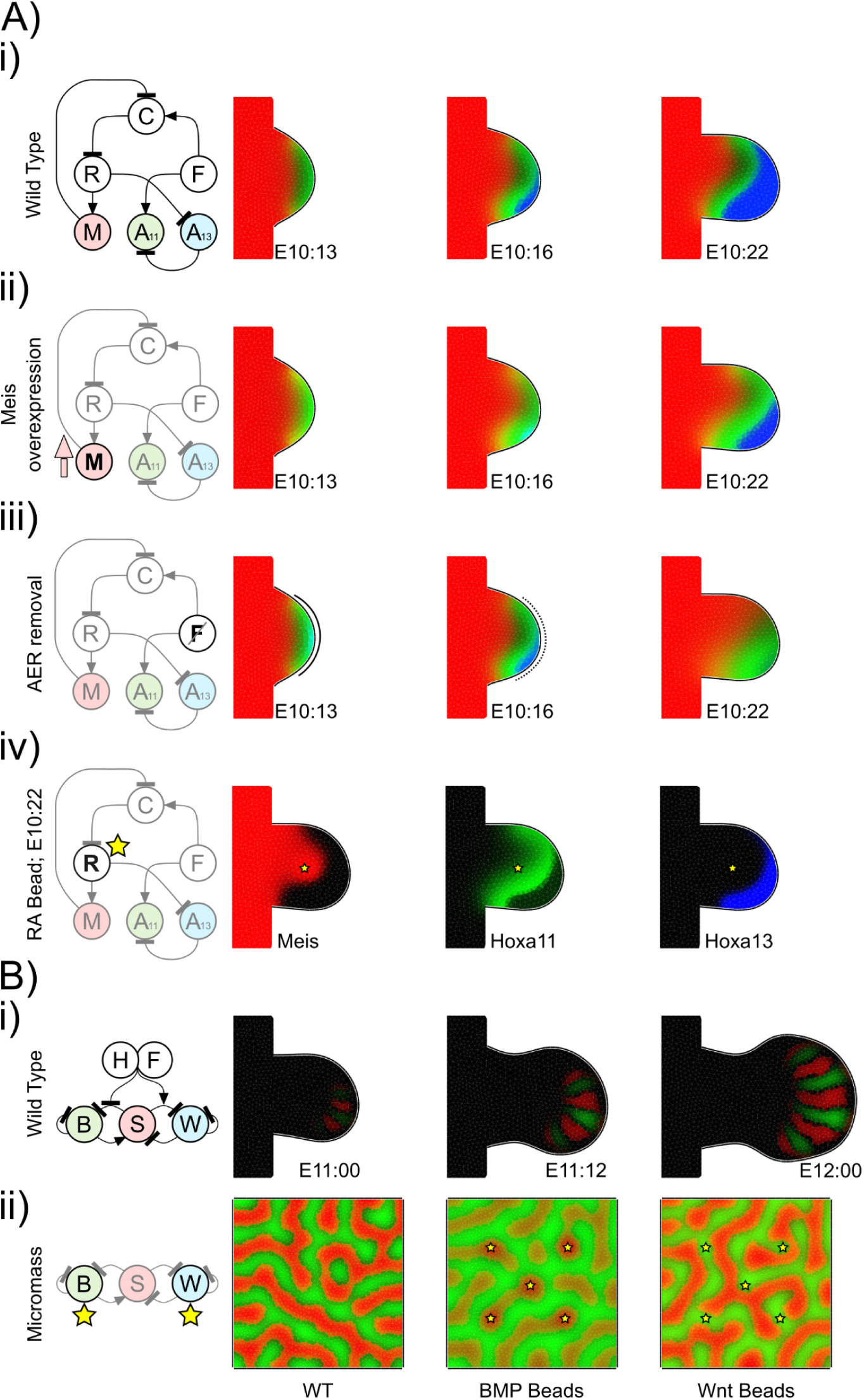
LimbNET can reproduce existing models. An illustrative sample of the variety of patterning mechanisms and experimental interventions that can be modelled using LimbNET. (A) Proximo-Distal (PD) patterning, an implementation of “Model F”, the final regulatory network scheme introduced by Uzkudun et al.(Uzkudun *et al*, 2015). Multiple experiments are shown via adjustments to the same model, illustrating the flexibility of LimbNET. (i) The “wild-type” model shows typically observed expression patterns of Meis (red), Hoxa11 (green) and Hoxa13 (blue). (ii) Global Meis overexpression induces a distal shift of the expression boundary between Hoxa11 and Hoxa13 (Mercader *et al*, 2009). (iii) Simulation of Apical Ectodermal Ridge (AER) removal at E10:16 (Vargesson *et al*, 2001). (iv) Introduction of beads soaked in retinoic acid induces localised overexpression of Meis and a distal shifting of the Hoxa11 pattern(Mercader *et al*, 2000). (B) The BSW model of digit patterning(Raspopovic *et al*, 2014), predicts correct number and initiation of digits following a Turing patterning mechanism involving a GRN of Bmp, Sox9 and Wnt signalling. (i) The “wild-type” patterning model simulated in a limb bud domain, reproduces the results of Raspopovic et al. Figure 3D; Sox9 shown in red, BMP shown in green. (ii) Further experimental interventions can be simulated: *left*, the network is simulated on a square domain, representing cell culture conditions; *centre*, we introduce BMP-soaked beads to the cell culture, which locally upregulate Sox9 expression; *right*, we introduce Wnt-soaked beads which locally repress Sox9 while upregulating its expression over a longer range.

To illustrate typical use of LimbNET, we started with this “wild-type” model - depicting experimentally observed behaviours of the system under normal conditions - and we built upon the original equations, incorporating a number of experimental interventions, as was done in the original paper(Uzkudun *et al*, 2015). Our unified model was expanded to create a single model specification that takes into account three experiments and their observed consequences. Firstly, global overexpression of Meis1 has been shown to induce a distal shift of the expression boundary between Hoxa11 and Hoxa13 (Mercader *et al*, 2009). This can be simulated by a uniform upregulation of Meis throughout the limb bud (Figure 4A, ii). Secondly, we simulated removal of the AER (Figure 4A, iii), which can cause a loss of expression of Hoxa13 (Vargesson *et al*, 2001). This simulation was easily implemented by adding a time-dependent expression into the equation of the production of FGF (Figure 4A, iii). Finally, implanting a bead soaked in retinoic acid into the middle of the limb bud has a similar impact as the Meis over-expression. It shifts distally the expression boundary between Hoxa11 and Hoxa13 (Mercader *et al*, 2000). A tool is provided within LimbNET allowing the user to define virtual bead experiments. Both the position and the timing of bead insertion can be defined, by clicking on a given triangle at a particular time-point. This definition can then be linked to the production of any variable in the model - in this case retinoic acid, resulting in a recapitulation of the originally published simulation (Figure 4A, iv).

The second model we chose is a very different type of GRN, whose spatial patterning dynamic is dependent on feedback loops. It is the proposed BSW model (Raspopovic *et al*, 2014), a Turing system to explain the patterning of digits through the interactions of Bmp, Sox9 and Wnt signalling. Unlike their more straightforward mechanistic relatives, these equations contain a number of nonlinear terms, time-dependent changes in distal signalling, and constraints on some of the spatial variables. This model was again reimplemented as a single LimbNET model specification, which upon simulation successfully reproduces the published results: the generation of a five-digit striped pattern in the limb bud autopod (Figure 4B, i), arising from a Turing patterning mechanism modulated by a time-dependent distal-to-proximal gradient of FGF. Secondly, we investigated the behaviour of the model under conditions more akin to a mesenchymal cell culture or micromass (Figure 4B, ii), by re-simulating the model specification in a square-shaped 2D domain. Through our model, we successfully reproduced experimentally observed results(Raspopovic *et al*, 2014): implanting virtual beads, soaked in either BMP or Wnt ligands, into virtual micromass culture results in local upregulation of sox9 in the case of BMP beads (Figure 4B, ii) and likewise in local inhibition and distal upregulation of sox9 expression for Wnt-soaked beads (Figure 4B, iii). Overall, LimbNET is able to successfully recapitulate previously published models and simulations, provided in a user-friendly web-based platform which allows users to explore, understand and modify the original models.

## Discussion

Understanding complex multi-scale biological phenomena will require computational modelling, but this will in turn require the sharing of models and simulations between a diverse community of theoreticians and experimentalists. It is important that predictive computer modelling facilitates collaboration and the shared exploration of new ideas and hypotheses. LimbNET is a first step towards this vision, specifically oriented towards the paradigmatic model system of limb development.

We believe that an open web-based simulation platform linked to a core common data framework provides the ideal scenario for getting scientists on the same page - ensuring both data and ideas can be compared in an objective way. Sharing data *per se* (Figure 5A) is only the first step. Two images of the same gene expression pattern from two different research groups may both be on the same website, and we may compare them by eye. It is true that the conclusions we wish to derive are often qualitative: “the mutant gene expression pattern was more proximal than the wildtype”. However, although that type of desired statement is qualitative, it requires a quantitative comparison to prove whether it is true; stepping forward from pure digitisation/quantification to objective comparison. While data and its analysis may often be used to *describe* a system, mathematical models represent hypotheses *explaining* the system, with the symbiosis between data and models cultivating new insights. By coupling our data framework to a computational modelling platform, we hope to help researchers more easily explain complex patterning data via dynamical modelling (Figure 5B). Simple causal relationships do not require computer modelling to explain or understand them. For example, the question “which upstream Transcription Factor activates gene X?” may be answered directly by a purely experimental approach, such as Chip-Seq(Osterwalder *et al*, 2014). Indeed a computational dynamical model cannot answer such a direct empirical question. However, if we want to understand how a molecular network is responsible for shaping dynamic spatial expression patterns over time, this is too complex to understand purely by mental thought processes - a true understanding can only be gained through model-based analysis of the system(Balaskas *et al*, 2012; Jaeger *et al*, 2004a).

**Figure 5:**
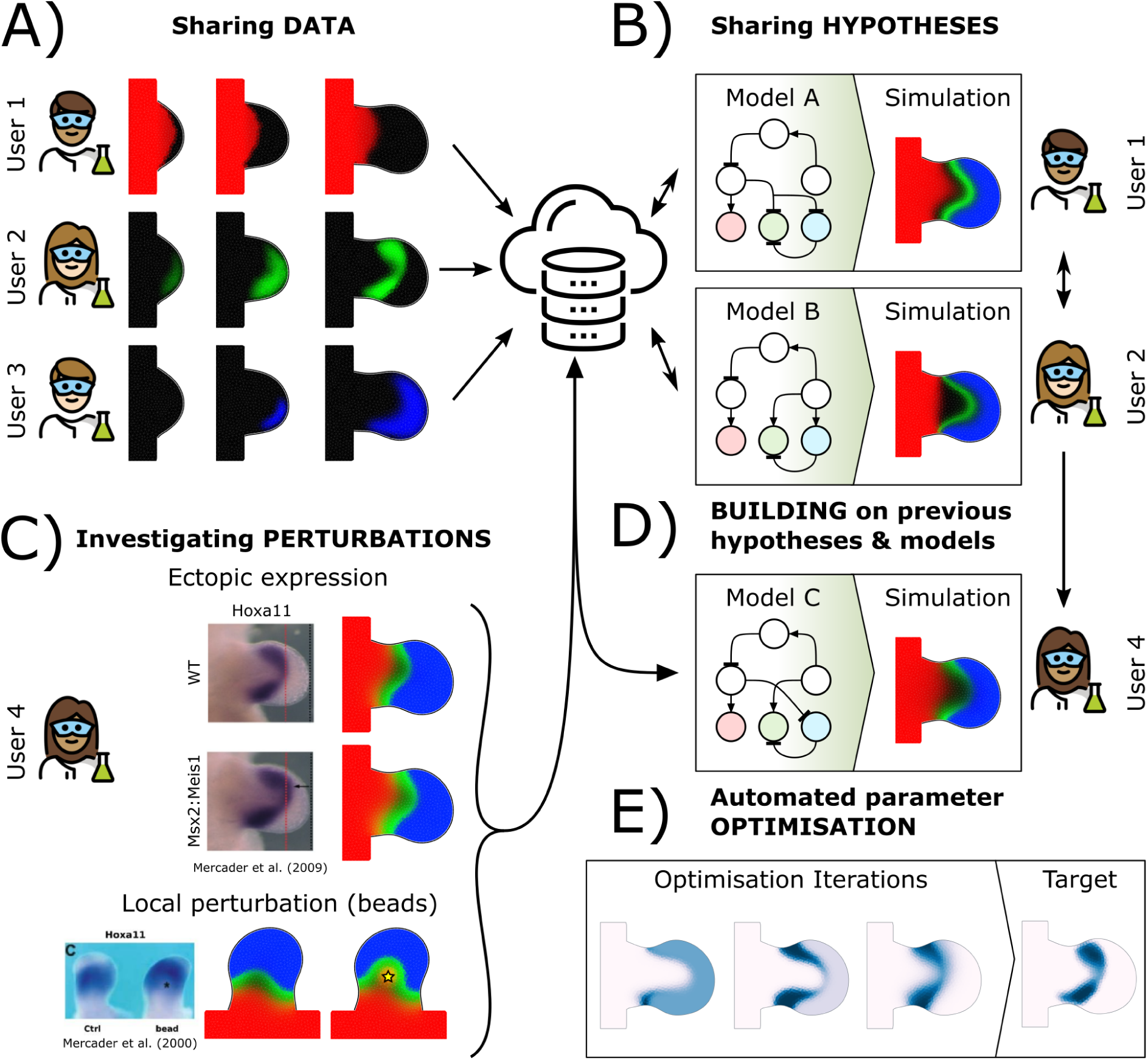
Hypothesis-driven open science. The most important novelty of LimbNET is the ability to share ideas/hypotheses plus the simulation tools to run them, rather than being purely a “database”. Nevertheless, its power lies in being data-driven, and it is indeed important that users are able to share spatiotemporal patterning data via a central data repository (A). In this illustrative example, three users have uploaded three separately acquired spatiotemporally varying gene expression patterns acquired from imaging data; Meis (red); Hoxa11 (green); Hoxd13 (blue). In addition to the expression data, LimbNET also provides for the sharing of hypotheses in the form of mathematical models (B). Users may upload models of regulatory networks specifying some form of spatiotemporally varying GRN dynamics, and simulate them. For example, two users observe the data uploaded in (A) and devise two different hypotheses for the underlying behaviour, formalised as two different regulatory network models (“Model A” and “Model B”). They upload the equations to LimbNET and are able to simulate them and compare the predictions generated by both interaction networks. Given a regulatory network, and corresponding mathematical model, users may compare the model output to further experiments (C), for example perturbations of the original experimental system. They - or indeed any other collaborators with access to LimbNET - may then refine or change the previous model(s) (D), essentially building on previous biological hypotheses and refining them to take into account new observations. (E) Automated parameter optimization is also an important tool within LimbNET. It is currently available through the API, but will soon have full access through the web-based client interface.

Although experiments are not sufficient to understand complex networks, data from experimental perturbations are nevertheless important in building, verifying or disputing model predictions, e.g. knock-outs, ectopic expression or beads experiments (Figure 5C). A single platform that includes both digitised experimental data and also model predictions greatly facilitates comparison of hypotheses with quantitative data (both for the wildtype case and experimental perturbations). Crucially this may lead to invalidation of model predictions under certain conditions (direct biochemical perturbations, mutations) thus offering further insights into hidden mechanisms, and the opportunity to (re)build existing work and foundations laid by prior modelling efforts (Figure 5D).

LimbNET’s tight integration between models/simulations and digitised data also allows researchers to automate parameter fitting of proposed regulatory networks to experimental observations (Figure 5E). This is currently available through the API, and shortly to be available in the GUI. Given a topology of regulatory network interactions, formalised as a system of PDEs, the user may choose certain sets of parameters to be automatically optimised to give the closest result to the empirical data. Parameter optimization operates by defining an objective function that attempts to minimise the discrepancy between simulation outputs and digitised data, in order to attempt to fit a specific network model to a concrete set of observations(Uzkudun *et al*, 2015; Lobo & Levin, 2015; Mousavi & Lobo, 2024).

We believe that LimbNET offers a number of advantages over existing modelling platforms. Users do not need to perform complex installation procedures - likewise, developers should not need to support many different Operating Systems - and the speed of simulation does not depend on users’ own hardware. A centralised resource (for both data and modelling) helps to ensure consistency of data (images, digitisations, models and simulations). Additionally this type of service facilitates sharing and collaboration(Byrne *et al*, 2010). Users can make data private, if need be, and conversely published models can be made public.

Reproducibility in biological modelling is a well-known challenge(Tiwari *et al*, 2021), where predominant issues include missing parameter values, initial conditions or inconsistent model structure. LimbNET dramatically reduces these problems. Although the backend (simulation engine) is remote/centralised, a model’s equations, parameters, etc., are all explicitly specified and open. A visiting researcher can immediately simulate a published result and verify it themselves. Crucially the visitor can also interrogate these published models through their own perturbations, modifying parameters and extending the existing equations.

Two features of LimbNET may be seen as limitations, but in fact we believe they strengthen the value of this platform to the community. Firstly, we wish the system to be both sophisticated in what it can achieve (a tight integration of both data analysis and predictive simulations) while also being very accessible, user-friendly and easy for non-specialists to use. To achieve this balance LimbNET is specifically focused on just one developmental organ (the limb bud) and on just one aspect of its development (the control of gene expression patterns by GRNs). More generalised software such as Chaste(Mirams *et al*, 2013) and Morpheus(Starruß *et al*, 2014) represent more general-purpose tools, configurable to simulate a much wider range of developmental systems and phenomena. However, the price for generality is a higher energy-barrier to non-specialist users. The goal for LimbNET is to focus on a particular community (limb development), and to make data-driven modelling as accessible as possible. Notwithstanding this goal, the LimbNET platform could indeed model a wide range of tissues and organs, if they can be represented by new 2D growing meshes.

Secondly, while LimbNET’s predefined wildtype tissue movements may at first appear limiting in terms of the limb developmental phenotypes that can be modelled, the current focus of LimbNET is to address the complexities of pattern formation by GRNs *per se*. By focusing on patterning rather than morphogenesis, LimbNET derives a major technical benefit in the speed of the computations, and therefore the usability of the system for researchers. In the near future we plan to add extra mesh trajectories that will represent different degrees of growth impairment, or excessive growth (such as the *Xt* mutant (Johnson, 1967), a mutation in the *Gli3* gene(Schimmang *et al*, 1992)). Nevertheless, it is important to note that a large proportion of reported genetic experiments (both knock-outs and ectopic expression) reveals shifts in gene expression patterns without significant growth defects. The majority of these have not yet been explained through GRN models of spatiotemporal signalling, something which will now be possible.

## Author Contributions

AM - Data curation, Formal analysis, Investigation, Methodology, Software development, Validation, Presentation, Writing JS - Conceptualization, Funding acquisition, Methodology, Project administration, Resources, Supervision, Presentation, Writing - review & editing

## Acknowledgements

We would like to thank EMBL’s IT hardware and support staff, in particular Thomas von Kiedrowski and BioIT. We thank members of the Sharpe Lab for useful discussions and feedback, especially Philipp Germann, Xavier Diego, Marco Musy, Heura Cardona, June JuYeon Han, Lau Avinyo and James Cotterell. This project was supported by funding from EMBL, an ERC Advanced grant (SIMBIONT, project no. 670555), and Spanish Plan Estatal Grants (PID2019-110868GB-I00 and BFU2015-68725-P).

## Footnotes

1 https://vuejs.org/

2 https://flask.palletsprojects.com

3 https://www.postgresql.org

4 https://redis.io/

5 https://python-rq.org/

